# Division of Labor in Hand Usage Is Associated with Higher Hand Performance in Free-Ranging Bonnet Macaques, *Macaca radiata*

**DOI:** 10.1101/011916

**Authors:** Mangalam Madhur, Desai Nisarg, Singh Mewa

## Abstract

A practical approach to understanding lateral asymmetries in body, brain, and cognition would be to examine the performance advantages/disadvantages associated with the corresponding functions and behavior. In the present study, we examined whether the division of labor in hand usage, marked by the preferential usage of the two hands across manual operations requiring maneuvering in three-dimensional space (e.g., reaching for food, grooming, and hitting an opponent) and those requiring physical strength (e.g., climbing), as described by Mangalam et al. [1], is associated with higher hand performance in free-ranging bonnet macaques, *Macaca radiata*. We determined the extent to which (a) the macaques exhibit laterality in hand usage in an experimental unimanual and a bimanual food-reaching task, and (b) manual laterality is associated with hand performance in an experimental hand-performance-differentiation task. We found strong negative relationships between (a) the performance of the preferred hand in the hand-performance-differentiation task (measured as the latency in food extraction; lower latency = higher performance), the preferred hand determined using the bimanual food-reaching task, and the normalized difference in the performance between the two hands (measured as the difference in the latency in food extraction between them normalized by the latency in food extraction using the preferred hand), and (b) the normalized difference in the performance between the two hands and the manual specialization (measured as the absolute difference in the laterality in hand usage between the unimanual and the bimanual food-reaching tasks; lesser difference = higher manual specialization). These observations demonstrate that the division of labor between the two hands is associated with higher hand performance.

## Introduction

Lateral asymmetries in body, brain, and cognition are almost ubiquitous among biological organisms [2-4]. An adaptationist would advocate that these asymmetries were evolutionarily selected because no bilateral organism can maneuver in three-dimensional space unless one side becomes dominant and always takes the lead [5]. Which side would become dominant, however, is beyond the scope of this hypothesis as there is no advantage or disadvantage evidently associated with either the left or the right side (see Glezer [6], an open-peer commentary on MacNeilage et al. [7]). Among all, manual asymmetries are a central theme of investigation because they are likely to have shaped primate evolution [8]. Manual asymmetries can manifest into (a) hand preference, that is, one hand majorly used while solving a unimanual task (which requires only one hand) or the hand used to execute the most complex action while solving a bimanual task (which requires both hands); (b) hand performance, that is, one hand used to execute actions more efficiently. Fagot and Vauclair [9] reviewed studies on individual-and population-level manual asymmetries among nonhuman primates and proposed the ‘task complexity’ theory which states that the extent of manual asymmetry increases with the complexity of the task (here, the complexity is defined by the spatiotemporal progression of the movements, i.e., coarse verses fine). Observations on several nonhuman primate species are consistent with the task complexity theory. For example, the relatively more complex bimanual food-reaching tasks have been found to elicit greater manual asymmetries than the unimanual versions of the same tasks in capuchin monkeys, *Sapajus* spp. [10,11] and *Cebus capucinus* [12], and chimpanzees, *Pan troglodytes* [13].

Besides exhibiting hand preference and hand performance, several nonhuman primates have also been found to exhibit manual specialization, that is, they preferentially use either the left or the right hand while solving some specific types of tasks. For example, while feeding arboreally, captive sifakas, *Propithecus* spp. preferentially used one hand to maintain postural support and the other hand to pluck leaves [14]. While extracting peanut butter from a PVC tube, wild Sichuan snub-nosed monkeys, *Rhinopithecus roxellana* [15], captive tufted capuchin monkeys [16], olive baboons, *Papio anubis* [17], and chimpanzees [18] preferentially used one hand to hold the tube and the other hand to extract the peanut butter. While foraging for scattered for the ground, captive gorillas, *Gorilla gorilla* [19] and chimpanzees [20] preferentially used one hand to take the food items towards the mouth, and the other hand to hold the remaining ones. While extracting peanuts from a lidded box captive tufted capuchin monkeys consistently used one hand to open the lid of the box and the other hand to reach for them [21]. While allogrooming, wild Sichuan snub-nosed monkeys [22] and both captive and wild chimpanzees [23] preferentially used one hand to hold the skin, and the other hand to remove dirt and ectoparasites. Mangalam et al. [1] argued that these observations might reflect specialization of the two hands for manual actions requiring different dexterity types (i.e., simple/complex hand movements in three-dimensional space, grasping, supporting the body, etc.), and along similar lines described division of labor in hand usage in free-ranging bonnet macaques, *Macaca radiata*. The macaques preferentially used the ‘preferred’ hand for manual actions requiring maneuvering in three-dimensional space (reaching for food, grooming, and hitting an opponent), and the ‘nonpreferred’ hand for those requiring physical strength (climbing). In a hand-performance-differentiation task that ergonomically forced the usage of one particular hand, the macaques extracted food faster with the maneuvering hand compared to the supporting hand, demonstrating the higher maneuvering dexterity of the maneuvering hand. However, whether such division of labor in hand usage improves hand performance in terms of the time and/or energy required to solve a given task remains unexplored.

In the present study, we examined whether the division of labor in hand usage, as described by Mangalam et al. [1], is associated with higher hand performance in free-ranging bonnet macaques, *Macaca radiata*. To this end, we determined the extent to which (a) the macaques exhibit laterality in hand usage in two experimental unimanual and a bimanual food-reaching task, and (b) manual laterality is associated with hand performance in an experimental hand-performance-differentiation task. We expected negative correlations between (a) the performance of the preferred hand in the hand-performance-differentiation task (measured as the latency in food extraction; lower latency = higher performance), the preferred hand determined using the bimanual food-reaching task, and the normalized difference in the performance between the two hands (measured as the difference in the latency in food extraction between them normalized by the latency in food extraction using the preferred hand), and (b) the normalized difference in the performance between the two hands and the manual specialization (measured as the absolute difference in the laterality in hand usage between the unimanual and the bimanual food-reaching tasks; lesser difference = higher manual specialization).

## Methods

### Subjects and Study Site

The subjects were 16 free-ranging bonnet macaques: 2 adult males – AM1 and AM2, 1 subadult male – SM1, 4 juvenile males – JM1, JM2, JM3, and JM4, 8 adult females – AF1, AF2, AF3, AF4, AF5, AF6, AF7, and AF8, and 1 juvenile female – JF1 (see Table 1), inhabiting the Chamundi Hill range in Mysore, India (GPS coordinates: 2°14'41"N 76°40'55"E). We provided the macaques with food-reaching tasks and observed the corresponding hand usage. We adhered to the American Society of Primatologists (ASP) “Principles for the Ethical Treatment of NonHuman Primates” and conducted the present study as a part of an ongoing research project that was approved by the Institutional Animal Ethics Committee (IAEC) at the University of Mysore (because we conducted our research on individuals which (a) did not belong to an endangered or a protected species, and (b) inhabited an unprotected land with an unrestricted public access, our research work did not require permission from any other authority).

**Table 1.**
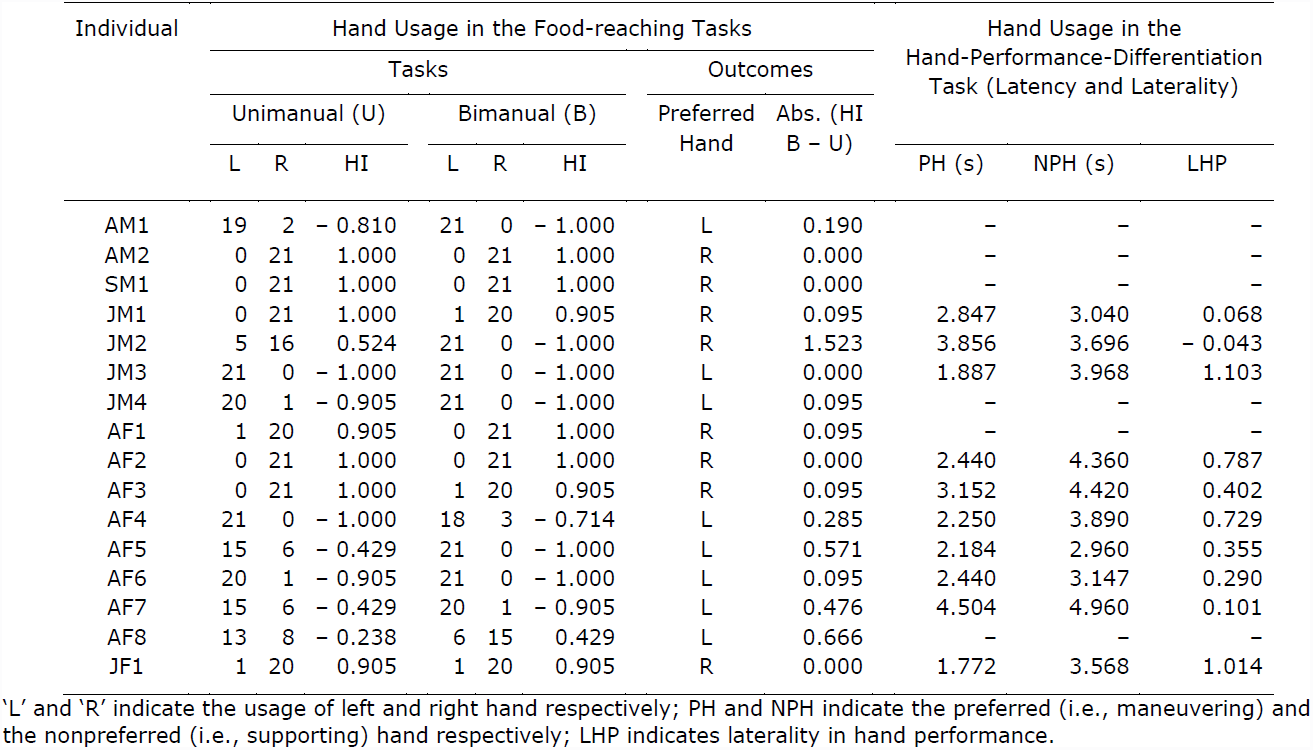
Raw data on hand usage for the macaques in the unimanual and the bimanual food-reaching tasks (n = 16), and the hand-performance-differentiation task (n = 10).

### Experimental Procedure

We presented the macaques with 3 sets of 7 consecutive trials, that is, 21 trials, of experimental unimanual and bimanual food-reaching tasks. Solving the unimanual task required obtaining a grape from an unlidded wire mesh box (dimensions: 7.5 cm × 7.5 cm × 17.5 cm; these dimensions allowed the usage of only one particular hand at a time) fixed on a wooden platform (dimensions: 90 cm × 60 cm) with one hand (Fig. 1A; Movie S1), whereas solving the bimanual task required opening and supporting the lid of a lidded wire mesh box with one hand and obtaining a grape with the other hand (Fig. 1B; Movie S2). We placed the task apparatus on the ground within ca. 1 m from the focal macaque when no conspecific was present within ca. 3 m from it and observed the corresponding hand usage.

**Figure 1.**
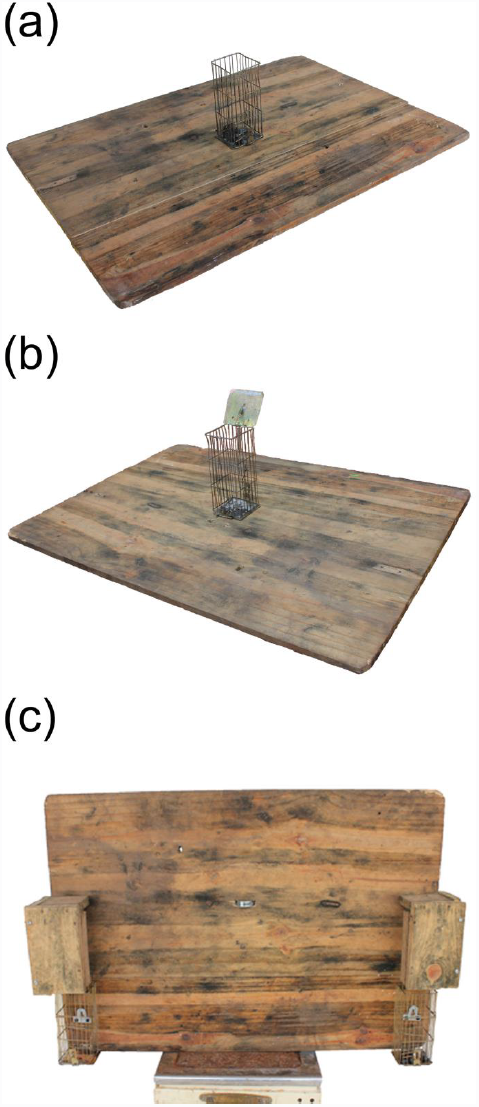
Apparatuses for the unimanual food-reaching task (a), the bimanual food-reaching task (b), and the hand-performance-differentiation task (c). Reproduced, with permission from Wiley Periodicals, Inc., from Mangalam et al. [1] © 2013 Wiley Periodicals, Inc.

We then presented the macaques with a single trial of an experimental hand-performance-differentiation task that forced the usage of either the left or the right hand. Solving this task required obtaining grapes from the wire mesh boxes attached towards the bottom on the either lateral extremities of a wooden platform (dimensions: 90 cm × 60 cm); this setup ergonomically forced the macaques to use either the left or the right hand (Fig. 1C; Movie S3). We put 7 grapes in one of the boxes, placed the task apparatus on the ground when no conspecific was present within ca. 3 m from the focal macaque, and video recorded the corresponding extraction behavior. We then repeated the same procedure, but this time by putting the grapes in the other box. The macaques mostly took 4 to 7 bouts to take all 7 grapes out of the box. We analyzed the obtained videos frame-by-frame to determine the average latency in food extraction for all the bouts (each bout measured from when the hand entered the box to when it exited) to the nearest 0.04 s.

For each macaque, we determined the handedness index (HI) values for taking the food out of the wire mesh box in the unimanual and the bimanual food-reaching tasks, using the formula: HI = (R − L)/(R + L) (where ‘R’ and ‘L’ represent the frequency of usage of the right and the left hand respectively). The obtained HI values ranged from − 1 to + 1, with positive values indicating a bias towards the right-hand use and negative values indicating a bias towards the left-hand use, and the absolute HI values indicating the strength of the bias. We then determined manual specialization using the formula: MS = abs. (HI bimanual – HI unimanual). We determined the hand majorly used for taking the food out of the box in the bimanual food-reaching task, which we referred to as the ‘preferred hand,’ and the opposite hand, which we referred to as the ‘nonpreferred hand’ (previously, in Mangalam et al. [1], we referred to these as the ‘maneuvering’ and the ‘supporting’ hand respectively). Moreover, we determined the laterality in hand performance (LHP) in the hand-performance-differentiation task, using the formula: LHP = (latency in food extraction using the nonpreferred hand – latency in food extraction using the preferred hand)/latency in food extraction using the preferred hand. The obtained LHP values ranged from − 1 to + 1, indicating the normalized difference in the performance between the two hands (w.r.t. the preferred hand).

### Statistical Analysis

We used the Spearman’s rank correlation test to determine the relationships between (a) the latency in food extraction using the preferred hand and the laterality in hand performance in the hand-performance-differentiation task, and (b) the LHP in the hand-performance-differentiation task and the difference in the HI values between the unimanual and the bimanual food-reaching tasks. Moreover, we used a Mann-Whitney U-test to make sure that there was no difference in the number of bouts between the two hands for taking all 7 grapes out of the box, which could have influenced these relationships.

## Results

Table 1 reports the raw data on hand usage for the macaques (whereas all 16 macaques responded to the unimanual and the bimanual food-reaching tasks, only 10 macaques responded to the hand-performance-differentiation task perhaps because of a lower motivation to solve a relatively more difficult and time-consuming activity). We found strong negative correlations between (a) the latency in food extraction using the preferred hand in the hand-performance-differentiation task and the laterality in hand performance (LHP) (r_s_ = − 0.772, n = 10, p = 0.020; Fig. 2A), and (b) the LHP in the hand-performance-differentiation task and the manual specialization (r_s_ = − 0.752, n = 10, p = 0.033; Fig. 2B). There was no difference between the two hands in the number of bouts for taking all 7 grapes out of the box in the hand-performance-differentiation task (U = 41.5, df = 9, p = 0.226).

**Figure 2.**
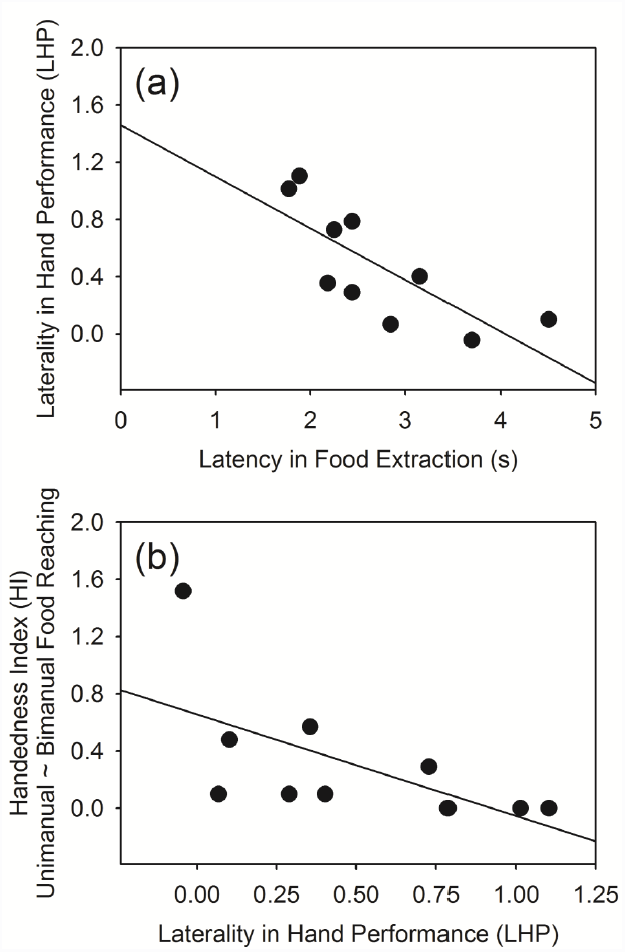
Relationship between the latency in food extraction using the preferred hand (i.e., the maneuvering hand, see Mangalam et al. [1]) and the laterality in hand performance (LHP) in the hand-performance-differentiation task (a), and the LHP in the hand-performance-differentiation task and the manual specialization (b). n = 10.

## Discussion

We examined whether the division of labor in hand usage, as described by Mangalam et al. [1], is associated with higher hand performance in free-ranging bonnet macaques. We found strong negative relationships between (a) the performance of the preferred hand in the hand-performance-differentiation task and the normalized difference in the performance between the two hands, and (b) the normalized difference in the performance between the two hands in the hand-performance-differentiation task and the manual specialization. These correlations demonstrate that the macaques that exhibit a higher manual specialization, show a greater difference in the performance associated with their two hands, and also extract food faster as compared to those that exhibit smaller differences.

On the one hand, the almost ubiquitous existence of manual asymmetries in nonhuman primates is likely to have some ecological advantages, and even more likely when there are underlying neurological asymmetries, as demonstrated in capuchin monkeys [24-27] and chimpanzees [28-30]. On the other hand, there may be some obvious disadvantages. Objects supposedly are randomly located with respect to the sagittal plane of an individual (i.e., towards the left or towards the right); this introduces difficulty in solving some tasks for individuals having a bias for one particular side. Fagot and Vauclair [9] reviewed studies on manual asymmetries in nonhuman primates and drew a distinction between hand preference and manual specialization. According to them, hand preference refers to the consistent usage of one hand to solve familiar, relatively simple, and highly practiced tasks, and may not be necessarily accompanied by an improvement in hand performance. In contrast, manual specialization refers to the consistent usage of one hand to solve novel, relatively complex, and not-practiced tasks that require peculiar action patterns, and is necessarily accompanied by an improvement in hand performance. Moreover, individuals generally exhibit manual specialization only when the tasks involve cognitively demanding manual actions. Thus, there exists a marked difference between hand preference and manual specialization in terms of the resulting differences in the performance of the two hands, which is evidently visible while considering the forms and/or functions of manual asymmetries, as described by Mangalam et al. [1]. The difference in the HI values between the unimanual and the bimanual food-reaching tasks allowed us quantifying manual specialization as an entity separate from hand preference (which an individual is likely to show because of an inherent bias) and examining whether it is associated with a higher difference in the performance between the two hands.

In a previous study [31], captive capuchin monkeys exhibited a weak, but statistically nonsignificant, positive relationship between the strength of hand preference and the corresponding hand performance in a unimanual and a bimanual versions of the box task. The study acknowledged that the strength of hand preference could have affected the timing of the movements, and so the observed relationship. This was, however, not the case of the present study because the hand-performance-differentiation task ergonomically forced the macaques to use either the left or the right hand, which allowed measuring the hand performance independent of any ceiling effects, i.e., it was unlikely to prime any motor actions associated with the hand opposite to that of the intended one. It provided a standard setup, which could be more widely used to compare hand performance across individuals while minimizing the possibilities of confounding effects. We suggest the developm
ent of such standard and robust experimental setups which might help answering the prevailing questions on manual asymmetries in nonhuman primates.

## Acknowledgements

This study was supported by a Department of Science and Technology (DST), Government of India, Ramanna Fellowship to MS.

## Supplementary Material

**Movie S1.** This footage illustrates the adult female bonnet macaque – ‘AF5’, solving the unimaunal food-reaching task.

Reproduced, with permission from Wiley Periodicals, Inc., from Mangalam et al. [1] © 2013 Wiley Periodicals, Inc.

**Movie S2.** This footage illustrates the adult female bonnet macaque – ‘AF5’, solving the bimanual food-reaching task.

Reproduced, with permission from Wiley Periodicals, Inc., from Mangalam et al. [1] © 2013 Wiley Periodicals, Inc.

**Movie S3.** This footage illustrates the adult female bonnet macaque – ‘AF5’, solving the hand-performance-differentiation task.

Reproduced, with permission from Wiley Periodicals, Inc., from [Mangalam et al. [1]] © 2013 Wiley Periodicals, Inc.

